# Estimating Organ-to-Plasma Ratios in physiologically based PK modeling: a simplified approach for early drug discovery

**DOI:** 10.1101/2024.03.21.586167

**Authors:** Fabio Broccatelli, Radha Ramakrishnan

## Abstract

Physiologically based pharmacokinetics (PBPK) modeling is a valuable tool in drug development and candidate selection. However, its widespread adoption in early stages of drug discovery remains limited. This study proposes a novel simplified approach to estimate organ-to-plasma ratio (K_p_) values based on measured or estimated volume of distribution (VD_ss_) to eliminate the reliance on additional in vitro data such as blood to plasma partition, LogP and pKa, which may not be available for newly synthesized molecules. The study includes 11 simulations across 4 compounds and 5 species. When simulations were based on ideal VD_ss_ and clearance (CL) values determined through non-compartmental analysis (NCA), the observed average absolute fold error (AAFE) aligned with experimental variability in the in vivo experiments (AAFE=1.3). However, when in vitro to in vivo correlation approaches were used to predict VD_ss_ and CL starting from in vitro intrinsic CL, plasma protein binding and microsomal binding, the AAFE increased to 1.7, reflecting the additional error introduced by the use of in vitro data to determine PK properties. Overall, this work provides a starting point for the facile implementation of PBPK models to meet the needs of early drug discovery projects.

## Introduction

Physiologically based pharmacokinetics (PBPK) modeling is utilized in drug development and during clinical candidate selection to predict human pharmacokinetics (PK) and to assess drug-drug interaction (DDI) risks.^1, 2^ PBPK has the potential to accelerate compound progression during the lead optimization phases, providing a platform to integrate in vitro and in vivo pre-clinical data towards human dose optimization, thereby reducing reliance on pre-clinical experiments. Early use of PBPK to support structure-activity relationships (SAR) has been explored by several investigators; despite the promising results, widespread adoption at this stage remains limited.^3, 4^ This is in part due to the limited software availability, a consequence of the cost of licensing commercial software running at scale as well as complexity of implementing proprietary PBPK models. In addition, the amount of data required to leverage PBPK models is a limiting factor during early stages; the methods most frequently employed to estimate organ to plasma ratio (K_p_) values rely on in vitro parameters such as lipophilicity, pKa, plasma protein binding and blood to plasma ratio, that may not be available for newly synthesized molecules.^5, 6^ While these parameters can be estimated by machine learning models, the expected deviations from the ideal values may result in large disconnects in the in vivo parameters estimates.^7^ K_p_ values are necessary in PBPK models to relate blood and organ concentration, hence methodologies that simplify the estimate of these parameters may increase the application of PBPK during early phases of drug discovery. In this study, we propose a novel approach to estimate organ K_p_ based on measured or estimated volume of distribution (VD_ss_). We demonstrate the usefulness of this approach in predicting PK across species starting from either PK parameters derived from non-compartmental analysis (NCA) or from a limited set of in vitro data (plasma protein binding, microsome binding and in vitro metabolic turnover).

### Physiological parameters

Physiological parameters for mouse, rat, dog, monkey, and human were obtained from various sources^8, 9^ Blood flow values for humans were derived from Wendling et al., while additional blood flow values for rodents were obtained from Yau et al. ^9 10^ All other physiological parameters, including aggregated values for plasma, blood, and tissues, were obtained from Davies and Morris.^8^ In cases where physiological values were not available for preclinical species, allometry was used to derive them. Missing mouse data were derived allometrically from rat, while missing monkey and dog data were derived allometrically from human^11^. An allometric exponent of 1 was used in the equation for weight values (e.g., organ weight), and an exponent of 0.75 was used for flow values (e.g., organ blood flow). The following organ compartments were considered: adipose, bone, brain, gut, heart, kidney, liver, lung, muscle, pancreas, skin, spleen, stomach, and the rest of the body (RoB). Blood volume (V_p_) was calculated as:

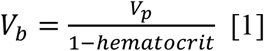

with a hematocrit value of 0.44 used across species.

The following body weight values were adopted: 0.02 kg for mouse, 0.25 kg for rat, 5 kg for monkey, 10 kg for dog, and 70 kg for human. The sum of all tissue volumes (V_t_) was normalized to match the V_t_ values as reported in Davies and Morris (0.557, 0.62, 0.51, 0.61, and 0.64 L/kg for human, mouse, rat, dog, and monkey, respectively). ^8, 12^

The blood flow for the hepatic artery was calculated as the difference between the liver and the splanchnic compartment (spleen, pancreas, stomach, and gut). For rat hepatic artery blood flow, corrections were made to the data from Yau et al. to ensure that the sum of the splanchnic compartment blood flow equated to the liver blood flow. ^10^ Data from Davies and Morris was used to estimate the blood flow via the hepatic artery in rats.^8^ This value was added to the liver blood flow and subtracted from the RoB blood flow. Dog, mouse and monkey blood flow for the RoB was not available; in dog this was calculated as a difference between total cardiac output and the sum of adipose, bone, brain, hearth, kidney, liver muscle and skin. In mouse and monkey, the same approach provided estimates that significantly differed from what was expected allometrically; hence allometry was used instead. This approach produced time constants (defined as ratio of blood flow and organ volume) within 2-fold across species for the RoB compartment. Table 1 includes all the normalized volumes and organ blood flow values.

**Table 1.** Normalized physiological parameters across different species.

|  | Mouse |  | Rat |  | Dog |  | Monkey |  | Human |  |
| --- | --- | --- | --- | --- | --- | --- | --- | --- | --- | --- |
|  | % Tissue | % Cardiac | % Tissue | % Cardiac | % Tissue | % Cardiac | % Tissue | % Cardiac | % Tissue | % Cardiac |

|  | Volume | Output | Volume | Output | Volume | Output | Volume | Output | Volume | Output |
| --- | --- | --- | --- | --- | --- | --- | --- | --- | --- | --- |
| Adipose | 7.6 | 10.9 | 9.1 | 7.0 | 20.9 | 2.9 | 21.5 | 2.5 | 27.8 | 5.0 |
| Bone | 2.1 | 19.8 | 2.5 | 12.2 | 7.0 | 5.7 | 7.2 | 8.9 | 8.6 | 5.0 |
| Brain | 0.5 | 3.1 | 0.6 | 2.0 | 0.7 | 3.8 | 1.9 | 12.0 | 2.3 | 12.0 |
| Gut | 7.2 | 15.9 | 2.5 | 11.0 | 4.5 | 18.0 | 2.6 | 15.5 | 1.6 | 14.0 |
| Heart | 0.5 | 3.5 | 0.4 | 4.9 | 1.1 | 4.5 | 0.4 | 7.4 | 0.5 | 4.0 |
| Kidney | 1.6 | 16.3 | 0.8 | 14.1 | 0.6 | 18.0 | 0.6 | 17.1 | 0.5 | 19.0 |
| Liver | 6.2 | 22.5 | 3.9 | 17.8 | 4.5 | 25.8 | 2.6 | 27.0 | 2.9 | 25.5 |
| Lung | 0.5 | 100.0 | 0.6 | 100.0 | 1.1 | 100.0 | 0.7 | 100.0 | 0.9 | 100.0 |
| Muscle | 48.0 | 11.4 | 44.7 | 27.8 | 51.8 | 20.8 | 48.3 | 11.2 | 45.8 | 17.0 |
| Pancreas | 0.3 | 0.5 | 0.4 | 1.8 | 0.1 | 0.4 | 0.1 | 1.3 | 0.2 | 1.0 |
| Skin | 13.9 | 5.1 | 21.0 | 5.8 | 3.3 | 8.3 | 9.7 | 6.7 | 4.0 | 5.0 |
| Spleen | 0.5 | 1.1 | 0.2 | 1.0 | 0.3 | 0.4 | 0.2 | 2.6 | 0.3 | 3.0 |
| Stomach | 0.4 | 0.7 | 0.5 | 1.3 | 0.2 | 0.4 | 0.2 | 1.3 | 0.2 | 1.0 |
| RoB | 10.7 | 7.4 | 12.9 | 8.4 | 3.9 | 10.3 | 4.0 | 7.2 | 4.4 | 7.5 |

The values used for the cardiac output are described in Table 2. In addition, the allometric relationship between cardiac output (CO) and body weight (BDW) was derived to enable future development of the model towards population modeling (R^2^=0.99, equation 2):

**Table 2.** Experimentally observed and estimated blood flow for liver and total cardiac output based on equation 2.

|  | Estimated Cardiac Output<br>ml/min | Reported Cardiac Output<br>ml/min | Estimated liver blood flow<br>ml/min/kg | Reported liver blood flow<br>ml/min/kg |
| --- | --- | --- | --- | --- |
| Mouse (0.02 kg) | 9 | 8 | 2 | 2 |
| Rat (0.25 kg) | 69 | 74 | 12 | 14 |
| Dog (10 kg) | 1369 | 1200 | 353 | 309 |
| Monkey (5 kg) | 781 | 1086 | 211 | 218 |
| Human (70 kg) | 6620 | 5600 | 1688 | 1450 |

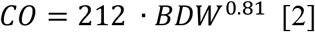

Blood flow for arterial and venous blood compartments were equal to CO, however the volume of the two compartment was different (venous 75% of the total blood volume, arterial 25% of the total blood volume).^9^

### PBPK Model Structure

Figure 1 describes the structure and the equations utilized in the whole body perfusion limited PBPK model presented in this study. Blood flow values for the respective organs are abbreviated as Q.

**Figure 1.**
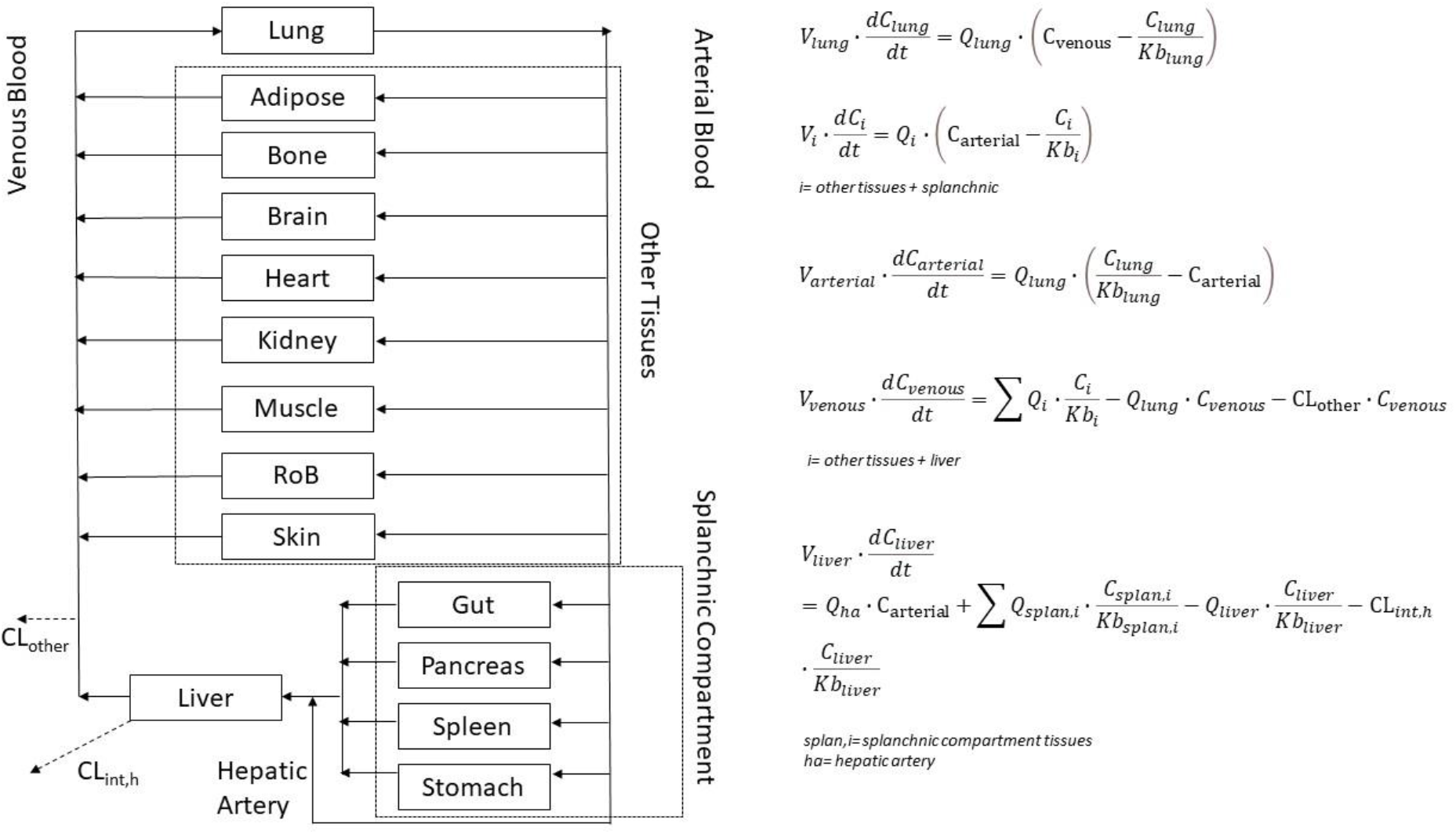
Structure of the PBPK model described in this work.

K_b_ and K_p_ are the tissue to blood and tissue to blood partition constants, respectively:

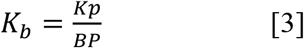

where BP is the blood to plasma concentration ratio.

The CL_int,h_ value represents the hepatic total intrinsic clearance, that is the product of fraction unbound in blood (fu_b_) and the free hepatic intrinsic clearance *CL*_*int,h,unbound*_ :

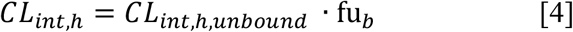

both f*u*_*b*_ *and CL*_*int,h,unbound*_ can be estimated from in vitro data.^13-15^ In addition, the hepatic CL_int,h_ can be back-calculated by leveraging the well stirred hepatic model^16^ from the total hepatic blood CL (CL_h_) under the assumption of complete hepatic elimination^16^:

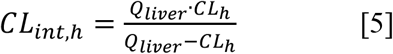

Fraction unbound in blood can be derived from fraction unbound in plasma:

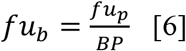

Equation 7 can be used to convert total hepatic plasma CL (CL_h,p_) into CL_h_ when PK parameters are estimated from plasma:

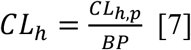

CL_other_ represents the non-hepatic clearance; this was represented as a composite parameter for multiple potential mechanisms in order to maintain a right-sized model structure for oral small molecule drug discovery.

The well-stirred model is utilized in this work for predicting blood or plasma CL (equations 8 and 9, respectively). The Korzekwa-Nagar is used for predicting VD_ss_ (equations 10-11)^16, 17^:

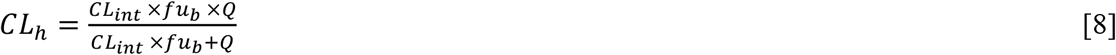

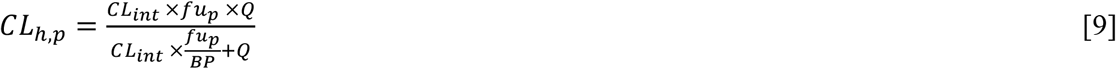

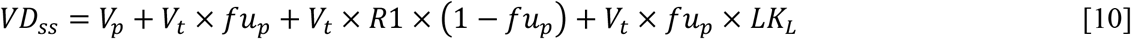

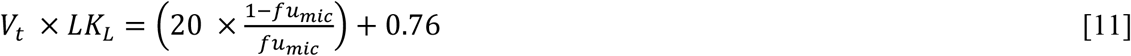

Where R1 is the concentration ratio of plasma proteins in tissue water-to-plasma, estimated to be between 0.052 and 0.116. An average value of 0.084 is adopted in this study. LK_L_ is the product of the tissue affinity and lipid concentrations for all the distribution compartment, this is estimated from fu_mic_. These PBPK models for PK parameter prediction are used in conjunction with the PBPK presented in figure 1 to predict the intravenous profile of a chemical entity starting from experimental measurements of fraction unbound in plasma (fu_p_) and microsomes (fu_mic_), as well as CL_int_ in microsome or hepatocytes.

### Derivation of plasma to tissue partition values

A collection of rat K_p_ values from various publications was obtained from Yun and Edginton.^18^ The data was used to establish a direct relationship between VD_ss_ and tissue K_p_. A total of 861 K_p_ values across 114 chemical entities and 10 tissues were available. The ionization classes from the original dataset were simplified into three categories: anionic, basic, and neutral. Trends for zwitterionic compounds were found to be consistent with the anionic compounds class, so these two categories were combined.

Predicted K_p_ values were derived leveraging a scalar to differentiate relative differences in K_p_ values between tissues (K_p,sim_) and a further K_p,scalar_ to equalize K_p,sim_ values based on VD_ss_.

Upon preliminary visual inspection, K_p_ values for most tissues correlated with either muscle or kidney ^5^ Adipose tissue was the only outlier. Visual analysis was conducted to assign K_p,sim_ values for each tissue within each ionization class based on correlations with K_p,muscle_ and K_p,kidney_. Median ratio between with K_p,kidney_ and K_p,muscle_ was found to be 3.8 across all compounds and ionization classes, hence K_p,sim_ values of either 3.8 (“kidney class”), 1 (“muscle class”) or 1.9 (“intermediate class”) were assigned to all tissues. These are summarized in in Table 3. In addition, plots comparing different tissues to muscle and kidney, as well as the tabulated VD_ss_ and K_p_ data, can be found in the supporting information. The correlation coefficient for the regression line and the number of predictions within twofold from identity were used as guidelines for the assignments. For tissues present in the PBPK model (Figure 1) but not available in the dataset used for the analysis, assignments were made based on similarity to muscle and kidney in terms of composition^5, 10^.

**Table 3.**
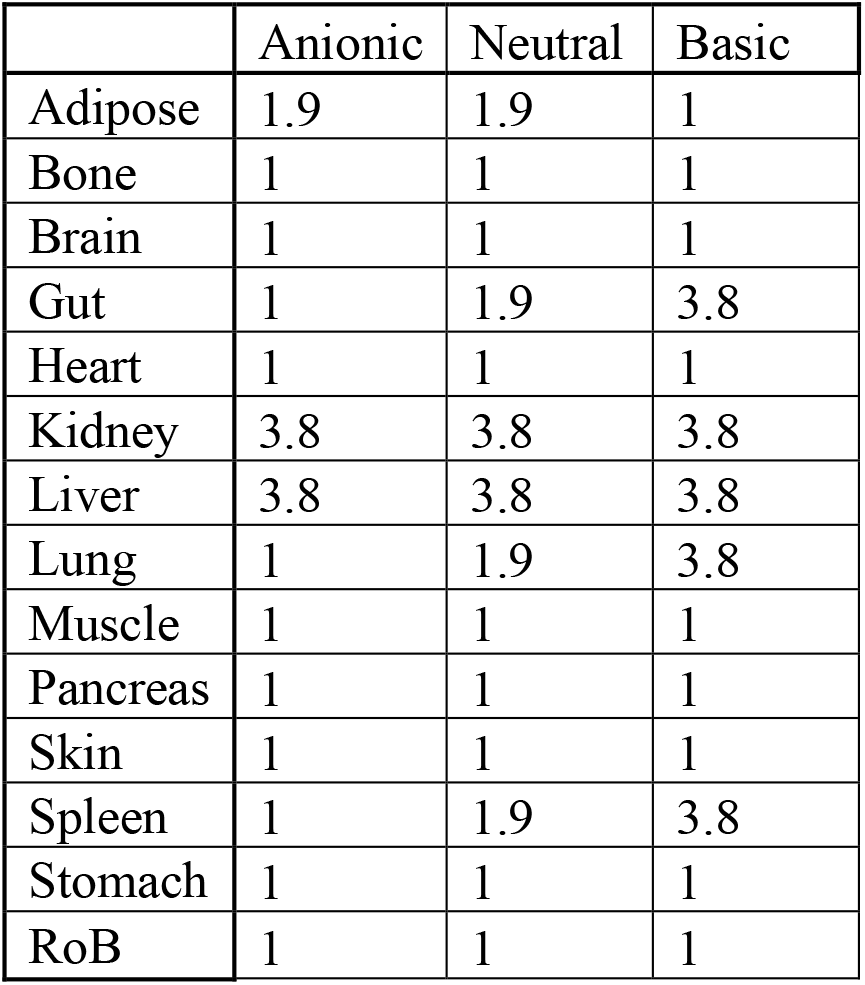
Relative K_p_ values based on tissue similarity (K_p,sim_)

Organ V_t_ values (V_i_) were obtained by multiplying V_t_ by the organ volume fraction as reported in table 1. The term K_p,scalar_, was derived by estimating the VD_ss_ in tissues (VD_ss,tissue_) from the experimental VDss, and by normalizing VD_ss,tissue_ by the theoretical VD_ss,tissue_ resulting from unequalized K_p,sim_ values (equations 8-10):

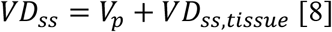

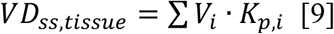

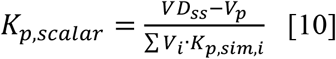

Finally, K_p_ for each tissue was calculated as:

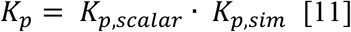

### Data analysis

Predictions were evaluated in terms of absolute average fold error (AAFE), root mean square error (RMSE) and % within 2, 3 and 5 fold from the experimental value:

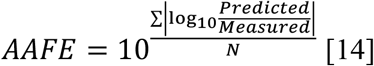

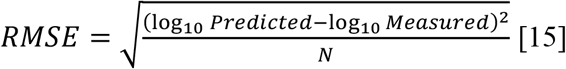

Where N represents the number of predictions.

### Comparison between PBPK derived and experimentally determined PK profiles

In vitro and in vivo data for verapamil, midazolam, warfarin and cefodizime was sourced from previous publications.^6, 13, 15, 19-25^ For midazolam, warfarin and verapamil, human PK profile was predicted starting from PK parameters determined by NCA analysis to test the ability to capture the shape of the PK profile starting from ideal PK parameters. In addition, prediction starting from PK parameters originated from in vitro data were also investigated.^16, 17^ For cefodizime the PK profile data was available in mouse, rat, dog, monkey and human; predictions of the PK curve across species starting from the ideal NCA PK parameters were produced to test the ability of the applicability of the model across different species.

## Results

### Plasma to tissue partition values prediction

Table 4 summarizes the performance of the algorithm presented in this work when predicting the experimentally determined rat K_p_ starting from VD_ss_:

**Table 4.** Accuracy of the K_p_ predictions method presented in this work. Value not reported in the referenced work are noted as NR.

|  | Count | AAFE | RMSE | % within 2fold | % within 3fold | % within 5fold |
| --- | --- | --- | --- | --- | --- | --- |
| **All Tissues** | 861 | 2.4 | 0.49 | 51% | 73% | 87% |
| **Adipose** | 68 | 3.8 | 0.70 | 28% | 43% | 62% |
| **Brain** | 90 | 3.1 | 0.64 | 42% | 60% | 73% |
| **All (excluded Adipose and Brain)** | 703 | 2.2 | 0.44 | 54% | 78% | 91% |
| **Muscle** | 107 | 2.2 | 0.42 | 50% | 81% | 91% |
| **All Tissues (Arundel)<sup>26</sup>** | 623 | 2.6 | 0.58 | NR | 69% | 82% |
| **All Tissues (Rodgers-Rowland)<sup>26</sup>** | 586 | 2.1 | 0.44 | NR | 77% | 88% |

Predictions for adipose tissues were significantly less accurate compared to all the other tissues (AAFE increased from 2.4 to 3.8). This is in line with reports from previous investigators, which led to the development of ad hoc models for the adipose tissue leveraging correlations with physico-chemical parameters.^5, 26, 27^ The same approach could be applied to the algorithm presented in this work, but is beyond the aim of this study. The analysis was further stratified by the brain tissue (for which the transporter activity is expected to skew K_p_ values), as well as muscle tissues, the most abundant tissue in human.^5, 28^ Higher AAFE (from 2.4 for all tissues to 3.1 for brain) was observed for predictions of brain K_p_, in line with the expected blood brain barrier transporters activity. While a direct comparison on the same dataset is not available, the performance of the model presented in this work appears to be slightly improved when compared to the alternative in vivo models requiring only VD_ss_ as input data for all tissues except adipose (Arundel method), and slightly less accurate when compared to in vitro tissues distribution models (Rodgers and Rowland method).^26^ The performance of the algorithm presented in this study is comparable to in vitro based tissue composition models when adipose and brain tissues are excluded.

### Prediction of PK profile in human

Figure 2 describes the performance of the PBPK model presented in this work in predicting the PK profile of Cefodizime across different species. The AAFE for the individual time-point predictions was 1.3 (1.6 for mouse, 1.3 for all the other species). The data suggest that the physiological parameters utilized across the different species are well calibrated.

**Figure 2.**
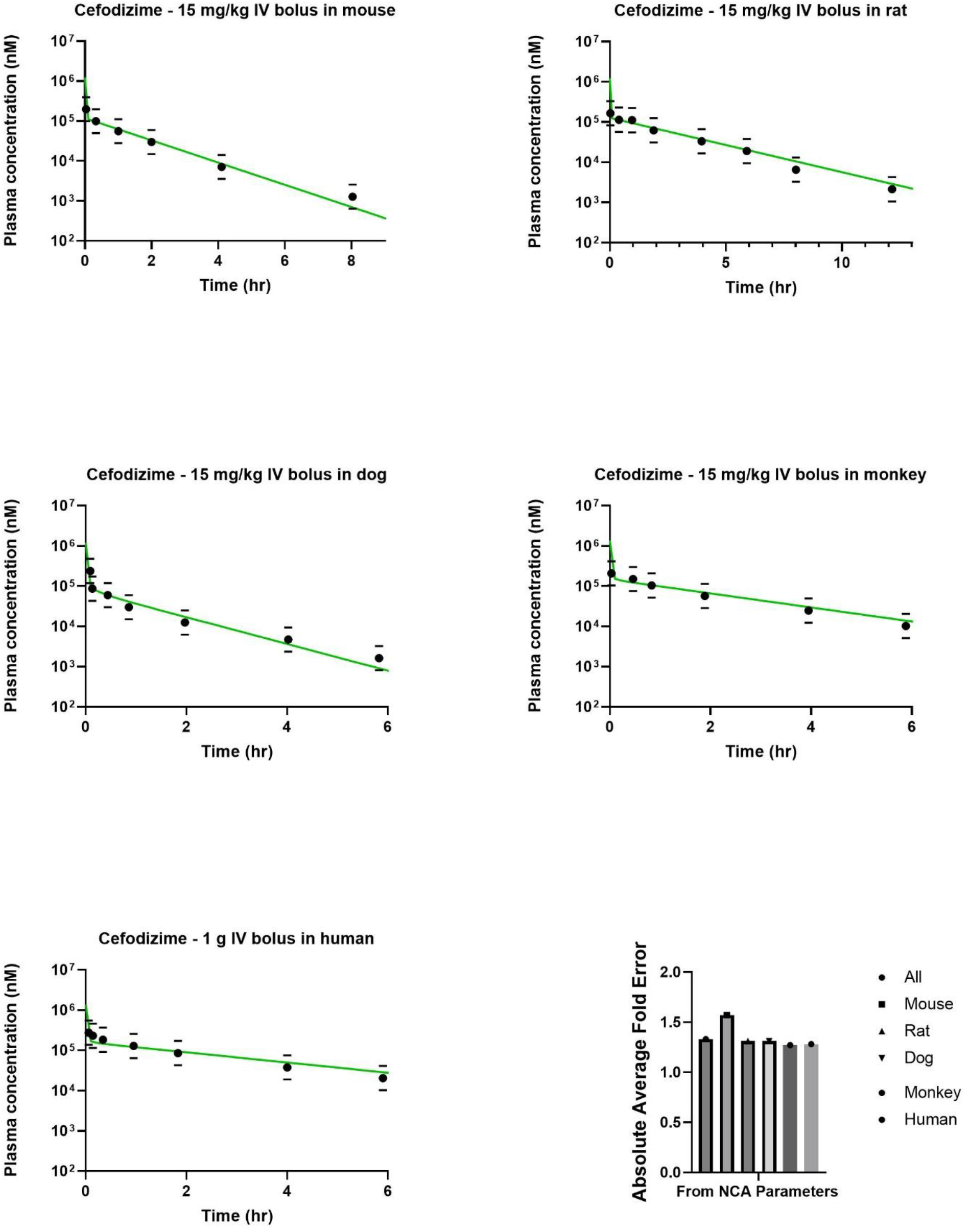
**PBPK simulation of experimentally determined plasma concentration - time profiles for cefodizime across five species. Green curves represent PBPK simulation resulting from the use of VD**_**ss**_ **and CL estimated from the experimentally observed PK profile via NCA. Black dots represent experimentally observed plasma concentrations, black horizontal lines represent 2-fold deviation boundaries for the experimentally observed plasma concentrations. Bar plots are used to quantify the absolute average fold deviation for the different PBPK simulations**.

Figure 3 extends the analysis to three prototypical Cytochrome P450 substrates in human. Similarly to what was seen for Cefodizime across species, an AAFE of 1.2 (1.3 for Verapamil,1.1 for Warfarin and Midazolam) was observed for the prediction of the individual time-points when the PBPK model was informed by the experimentally observed in vivo PK parameters from NCA. The AAFE increased to 1.7 when the PK parameters were predicted from the in vitro metabolic stability, plasma protein binding and microsome binding (1.9 for midazolam, 1.5 for warfarin, 1.4 for verapamil).

**Figure 3.**
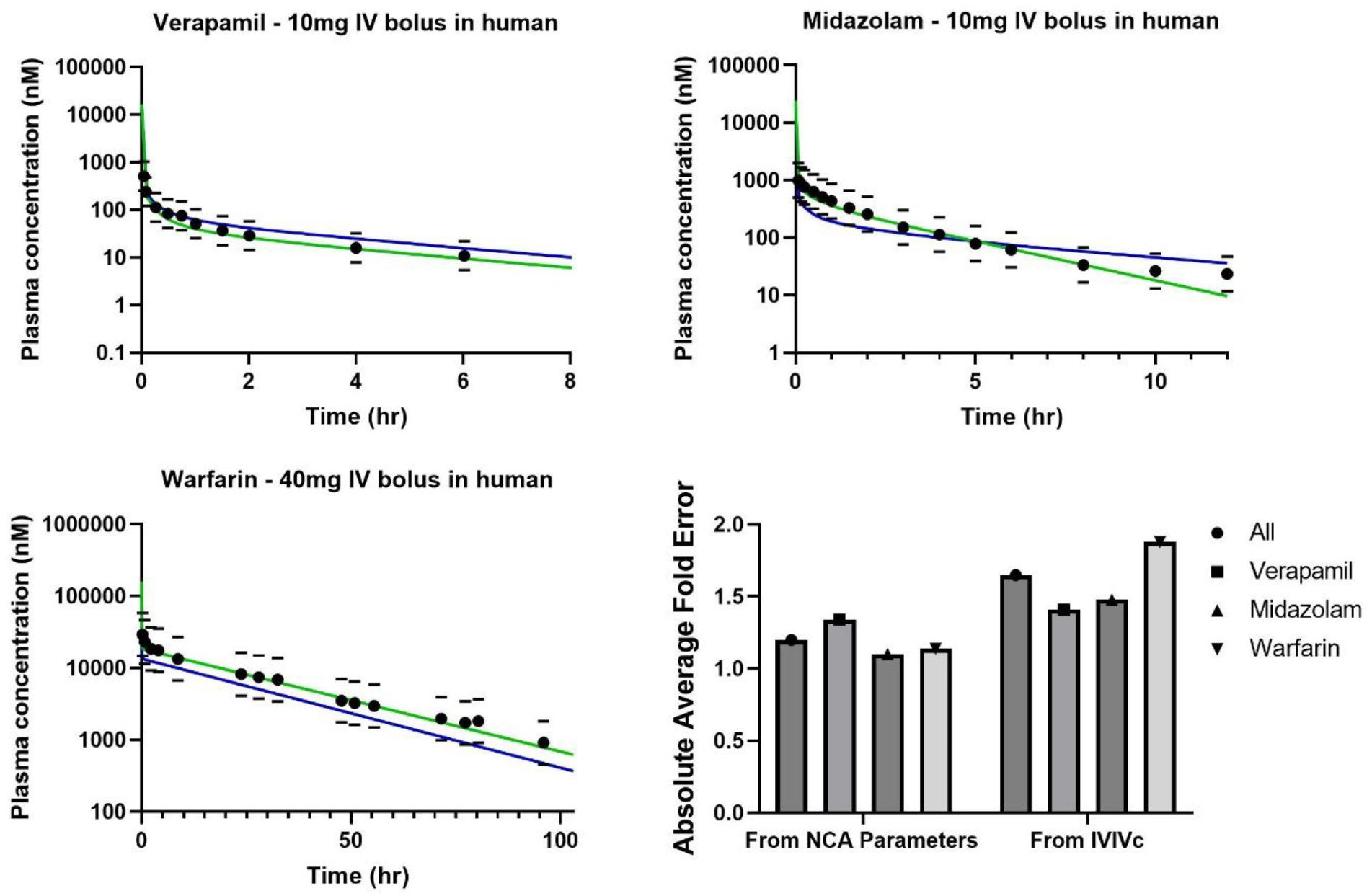
**PBPK simulation of experimentally determined plasma concentration - time profiles for verapamil, midazolam and warfarin in human. Green curves represent PBPK simulation resulting from the use of VD**_**ss**_ **and CL estimated from the experimentally observed PK profile via NCA. Blue curves represent PBPK simulation resulting from the use of VD**_**ss**_ **and CL estimated from in vitro data. Black dots represent experimentally observed plasma concentrations, black horizontal lines represent 2-fold deviation boundaries for the experimentally observed plasma concentrations. Bar plots are used to quantify the absolute average fold deviation for the different PBPK simulations**.

## Discussion and Conclusion

This work introduced a simplified approach to predicting organ K_p_ within the context of a perfusion-limited intravenous PBPK model for small molecules. The algorithm directly predicts K_p_ values from VD_ss_, eliminating the need for additional in vitro data. Although the accuracy of K_p_ prediction is slightly lower compared to in vitro based tissue partitioning approaches, this model does not rely on additional in vitro parameters that may not always be available during early phases of drug discovery (e.g. experimental LogP, pK_a_ and BP). Therefore, the model presented in this work is particularly suitable for early phases of drug discovery and lead optimization. Moreover, deriving K_p_ values directly from the input VD_ss_ allows for a clear separation between analyzing the shape of the PK profile (based on NCA PK parameters) and the accuracy of predicting PK parameters from in vitro data.

This study presented a total of 11 simulations involving 4 compounds and 5 species. When the simulations were based on the NCA VD_ss_ and CL values the observed AAFE was 1.3, which aligns with the experimental variability observed in in vivo experiments. This suggests that the PBPK model effectively captures the shape of PK profiles across different compounds and species, based on the dataset presented in this study.

However, when the model input relied on in vitro to in vivo correlation approaches, specifically the well-stirred hepatic model and the Korzekwa-Nagar model for VD_ss_, the AAFE increased to 1.7. This increase reflects the additional error introduced by the in vitro to in vivo correlation (IVIVc) approach. It is important to note that the compounds used in the IVIVc analysis are well-behaved and extensively characterized prototypical CYP450 substrates. Compounds emerging from drug discovery campaigns may have larger unknowns associated, leading to larger IVIVc disconnects.

All the physiological parameters and equations used in model building are provided in the manuscript, and the simulations and experimental PK profiles are included as supporting information. The simulations are generated using a minimal number of measured input parameters that are routinely generated in drug discovery projects in early PK assay tiers.

Overall, this work serves as a starting point for the easy implementation of high-throughput PBPK models to meet the requirements of early drug discovery projects.

