## Supplemental Figure 1 for "Estimating Organ-to-Plasma Ratios in physiologically based PK modeling: a simplified approach for early drug discovery"

**Tissue Kp correlations across different ionization classes (A=Anionic, B= Basic, N= Neutral, Z= Zwitterionic)**

Spleen vs. Muscle

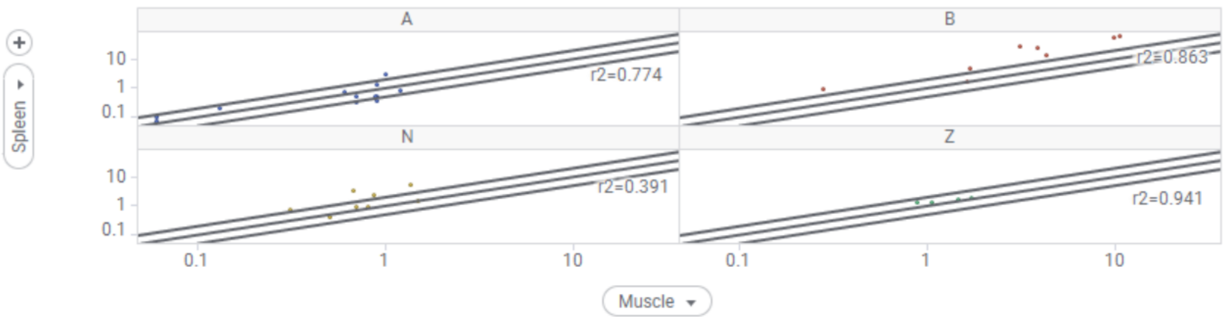


Spleen vs. Kidney

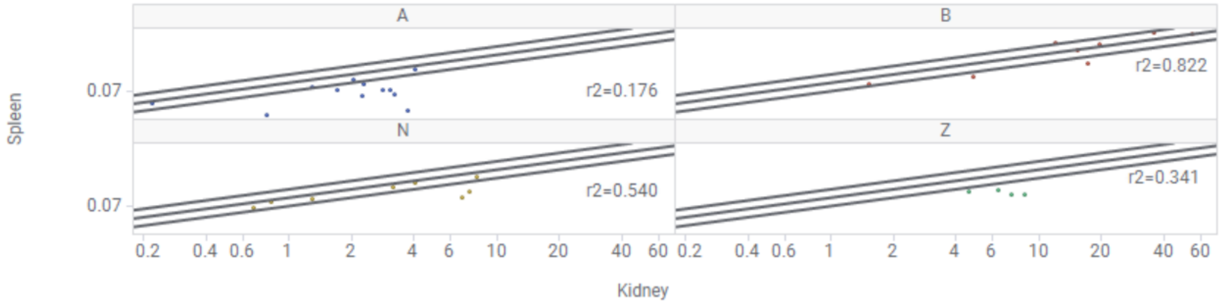


Skin vs. Muscle

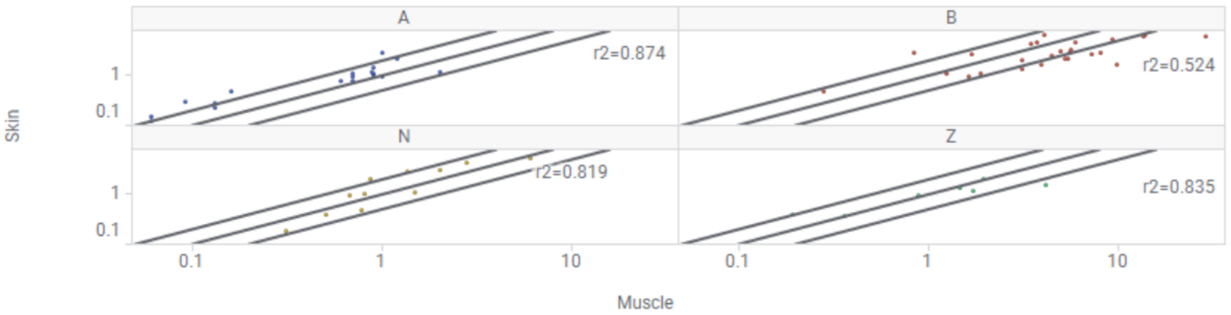


Skin vs. Kidney

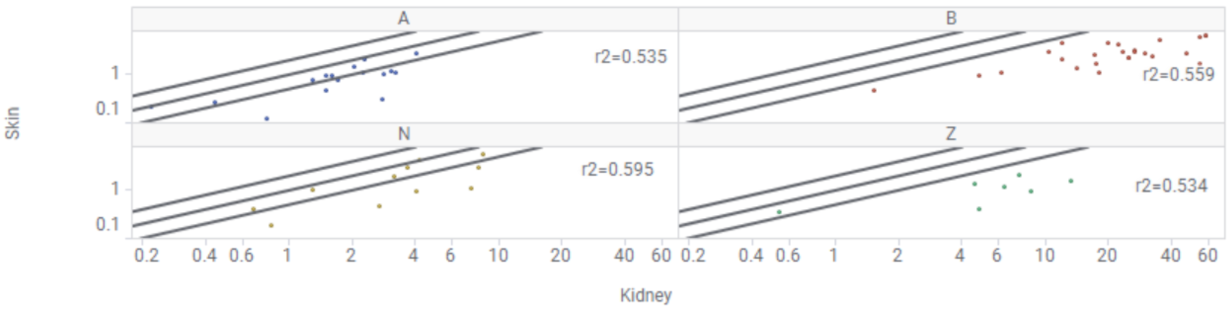

### Lung vs. Muscle

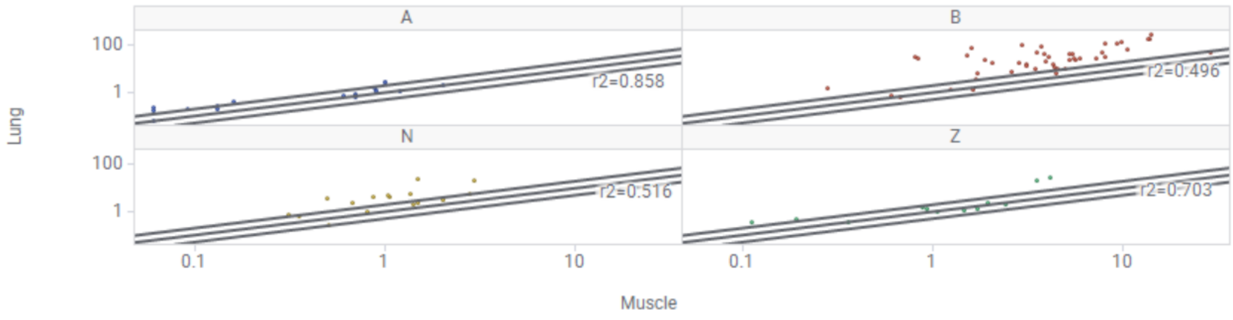


### Lung vs. Kidney

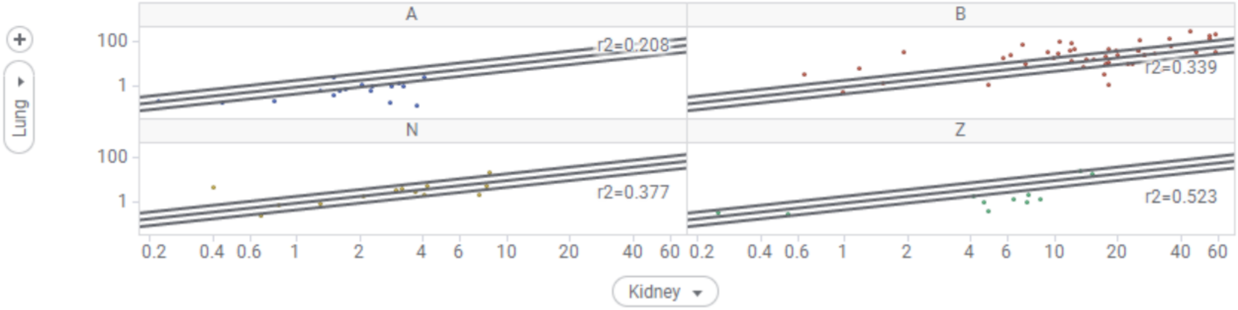


### Liver vs. Muscle

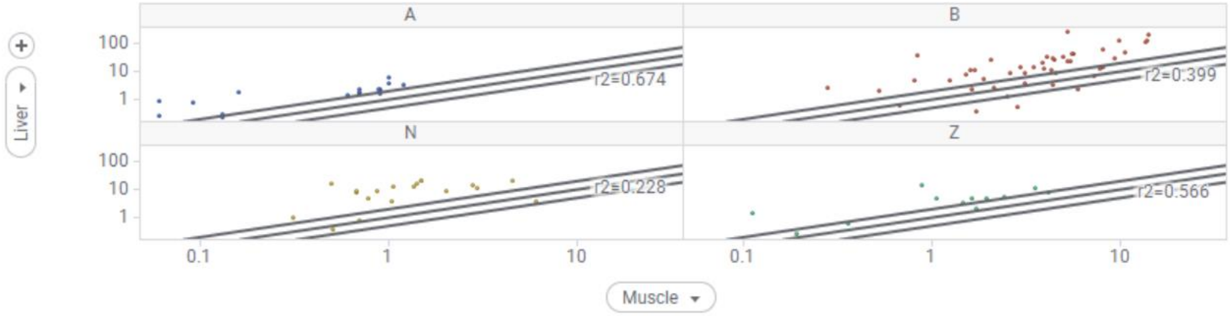


### Liver vs. Kidney

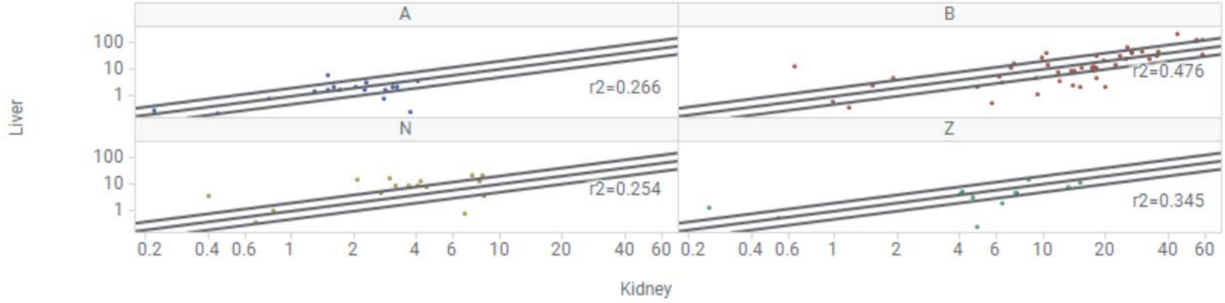

### Heart vs. Muscle

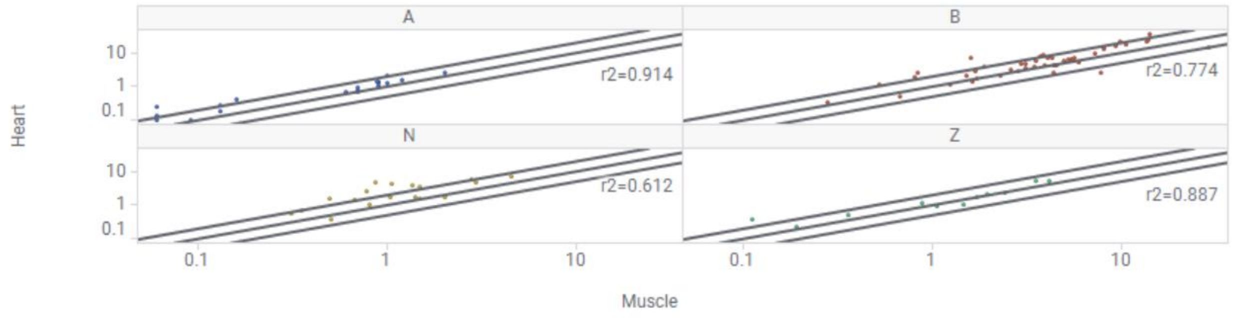


### Heart vs. Kidney

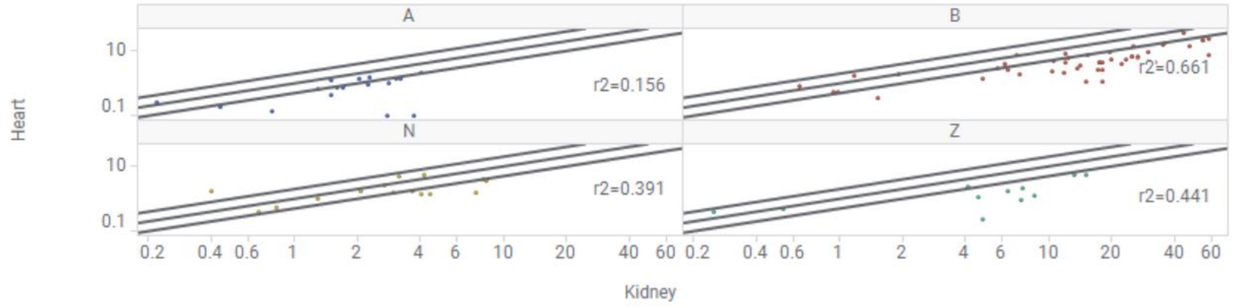


### Gut vs. Muscle

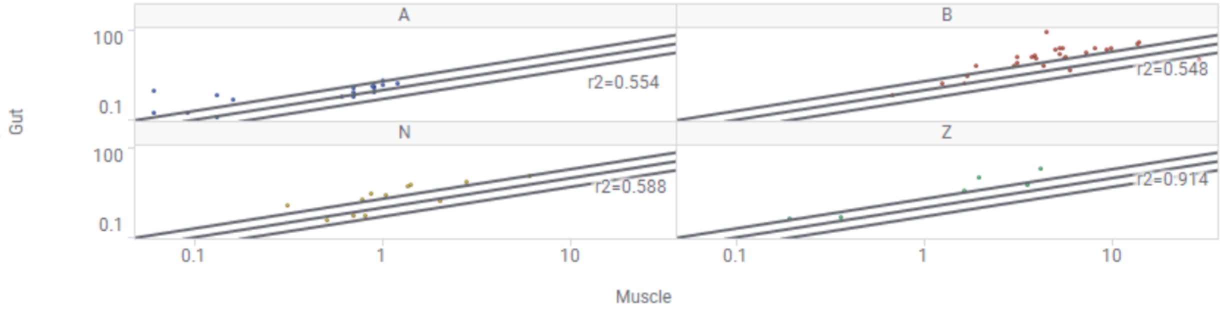


### Gut vs. Kidney

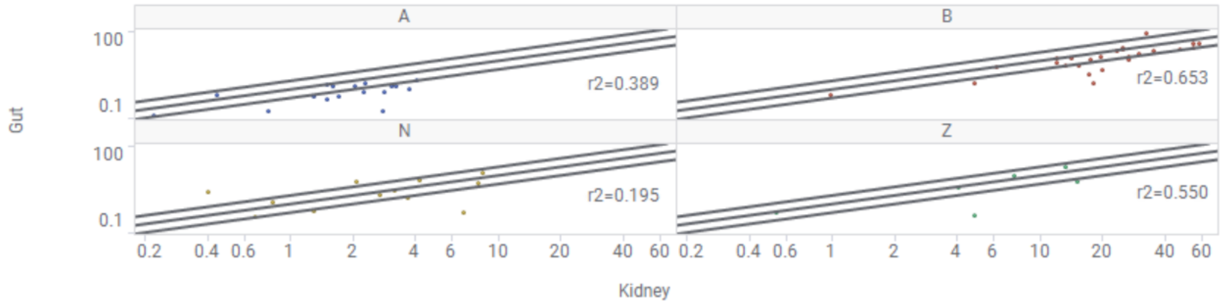

### Brain vs. Muscle

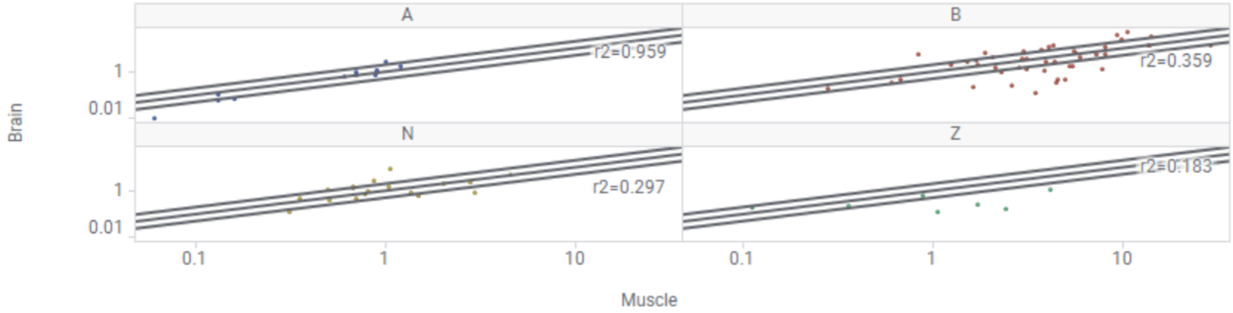


### Brain vs. Kidney

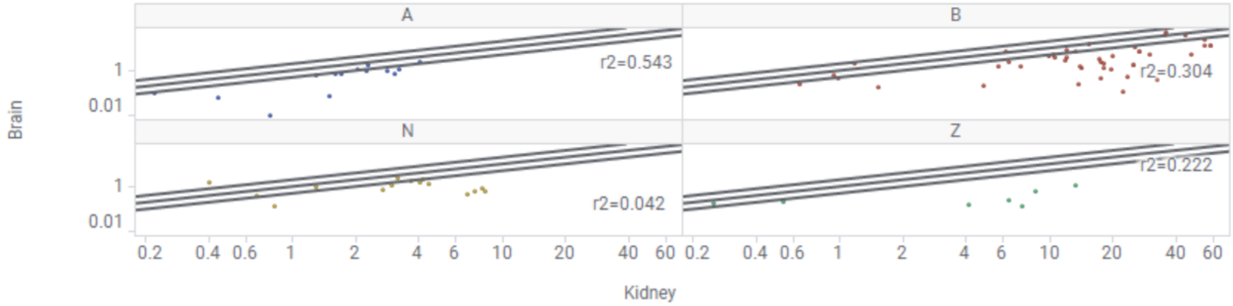


### Bone vs. Muscle

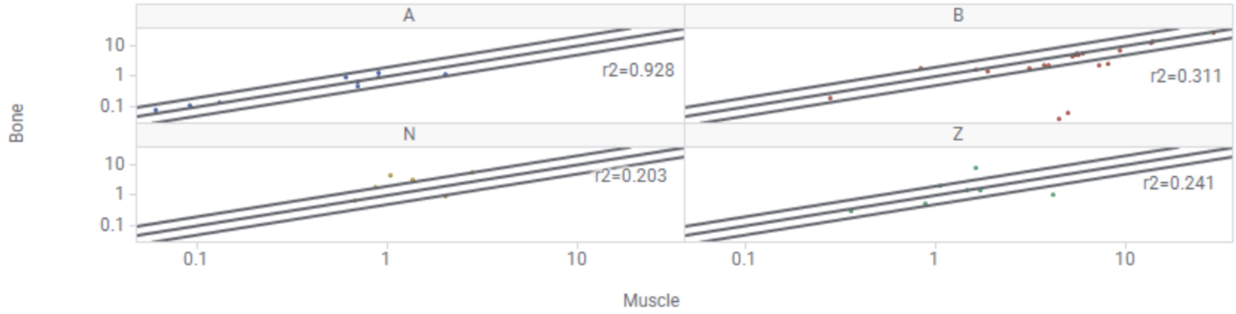


### Bone vs. Kidney

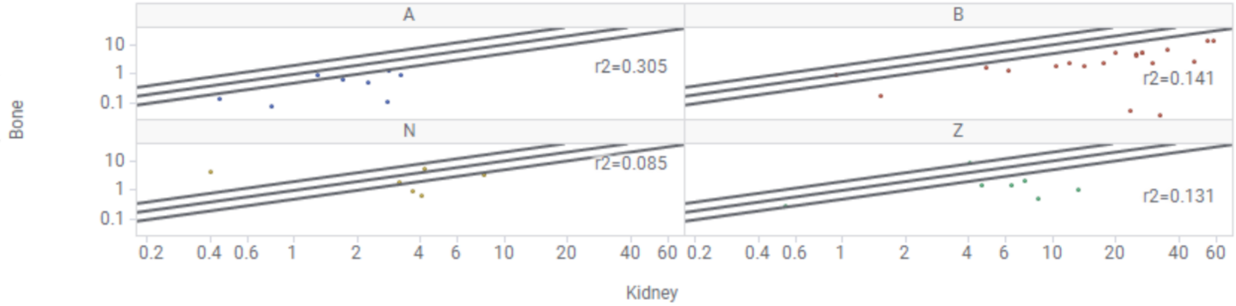

### Adipose vs. Muscle

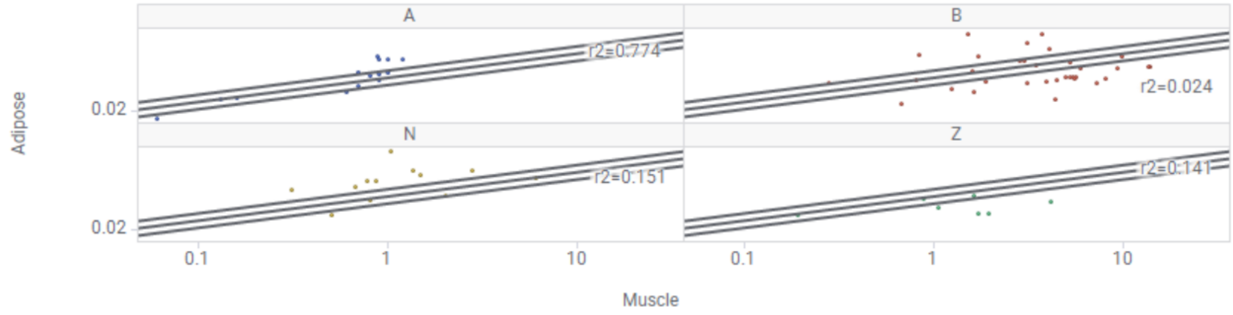


### Adipose vs. Kidney

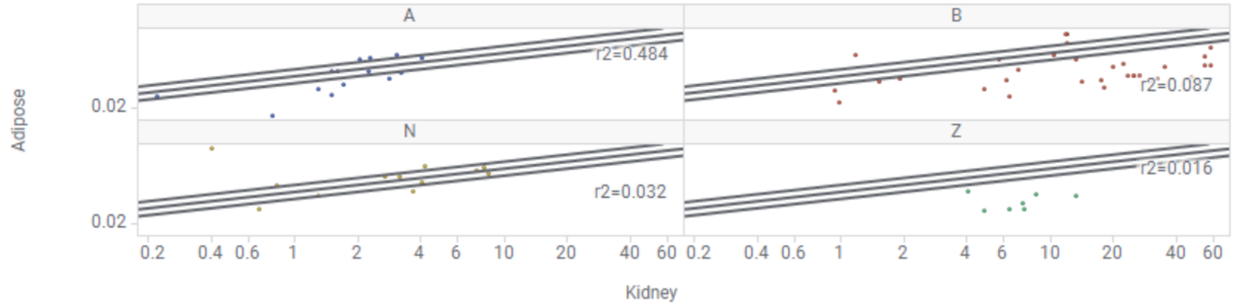
